# Synapsin condensation is governed by sequence-encoded molecular grammars

**DOI:** 10.1101/2024.08.03.606464

**Authors:** Christian Hoffmann, Kiersten M. Ruff, Irina A. Edu, Min Kyung Shinn, Johannes V. Tromm, Matthew R. King, Avnika Pant, Hannes Ausserwöger, Jennifer R. Morgan, Tuomas P. J. Knowles, Rohit V. Pappu, Dragomir Milovanovic

## Abstract

Multiple biomolecular condensates coexist at the pre- and post-synapse to enable vesicle dynamics and controlled neurotransmitter release in the brain. In pre-synapses, intrinsically disordered regions (IDRs) of synaptic proteins are drivers of condensation that enable clustering of synaptic vesicles (SVs). Using computational analysis, we show that the IDRs of SV proteins feature evolutionarily conserved non-random compositional biases and sequence patterns. Synapsin-1 is essential for condensation of SVs, and its C-terminal IDR has been shown to be a key driver of condensation. Focusing on this IDR, we dissected the contributions of two conserved features namely the segregation of polar and proline residues along the linear sequence, and the compositional preference for arginine over lysine. Scrambling the blocks of polar and proline residues weakens the driving forces for forming micron-scale condensates. However, the extent of clustering in subsaturated solutions remains equivalent to that of the wild-type synapsin-1. In contrast, substituting arginine with lysine significantly weakens both the driving forces for condensation and the extent of clustering in subsaturated solutions. Co-expression of the scrambled variant of synapsin-1 with synaptophysin results in a gain-of-function phenotype in cells, whereas arginine to lysine substitutions eliminate condensation. We report an emergent consequence of synapsin-1 condensation, which is the generation of interphase pH gradients realized via differential partitioning of protons between coexisting phases. This pH gradient is likely to be directly relevant for vesicular ATPase functions and the loading of neurotransmitters. Our study highlights how conserved IDR grammars serve as drivers of synapsin-1 condensation.

**Research Highlights:**

- Distinct biomolecular condensates formed by pre-synaptic proteins contain a unique bias of sequence grammars within their IDRs.
- The IDR of synapsin-1, the essential protein for the clustering of synaptic vesicles, has a conserved compositional bias for Arg and blockiness of proline/polar residues.
- Using sequence designs, we uncovered how conserved sequence features of the driver IDR of synapsin-1 affect condensation in vitro and in cells.
- Synapsin-1 condensates are defined by a measurable interphase pH gradient.

## Introduction

Functional neurotransmission relies on repeated cycles of synaptic vesicle (SV) exocytosis and endocytosis at the synapse. As part of this cycle, hundreds of SVs are clustered at the pre-synapse, forming distinct biomolecular condensates driven by highly abundant synaptic proteins including synapsins [1] [2], [3]. Synapsin-SV condensates enable reversible sequestration and maintenance of soluble synaptic proteins that are essential for distinct steps of the SV cycle [4] [5]. Pre-synapses appear to be complex emulsions defined by multiple coexisting phases composed of synapsins, synaptophysin, synucleins, and other pre-synaptic proteins [5].

The C-terminal intrinsically disordered region (IDR) in synapsin-1 is required for driving phase separation in cells and in vitro in the presence of macromolecular crowders such as polyethylene glycol [1]. Additionally, antibody binding to this IDR disperses SV clusters at lamprey synapses [2]. In cells, co-expression of synapsin-1 with synaptophysin (SYP), a resident SV membrane protein, is required for condensation [6]. Interactions between the cytosolic tail of SYP and the C-terminal IDR of synapsin-1 drive condensation [6]. The working hypothesis is that complementary electrostatic interactions contribute to the co-condensation of SYP and synapsin-1 [6] [7] [8]. The synapsin-1 IDR can also interact with SH3 domain-containing proteins, including intersectin, to drive condensation in vitro and regulate nanoscale organization in cells [1] [9]. Taken together, the extant data suggest that both IDR-IDR and IDR-structured domain interactions are important for the formation and regulation of synapsin-SV condensates.

It is generally thought that IDRs engage in weak, multivalent, and promiscuous interactions [10] [11]. Extrapolations from this thinking lead to the inference that phase separation is driven purely by non-specific interactions [12]. However, several discoveries have upended this thinking [13-28]. IDRs engage in sequence-specific interactions, and the affinities can span a diverse range depending on the complexes in question [29] [30] [31] [32] [33]. The sequence specificity of interactions involving IDRs is evident in their conformational ensembles, functions, and phase behaviors [28] [34-54]. Efforts over the past fifteen years have yielded quantitative and decodable heuristics that explain the physico-chemical basis of IDR specificity [19] [48] [55] [56] [57]. From an evolutionary perspective, while the precise sequence of an IDR might be poorly conserved, alignment-free methods have helped uncover the conservation of non-random amino acid compositional biases and binary sequence patterns across orthologous and paralogous IDRs [48] [55]. These molecular grammars enable chemical specificity [58] [59] via the multivalence of sequence-encoded hierarchies of complementary interactions. Chemical specificity contributes to system-specific networks of multivalent homotypic and heterotypic IDR-mediated interactions [51] [52].

Here, we report that pre-synaptic proteins feature IDRs with evolutionarily conserved molecular grammars that are distinctive from the underlying IDRome and are suggestive of complementary interactions within the network of IDRs that make up SVs. Mutations to the synapsin-1 IDR that perturb its conserved molecular grammars can abolish SV condensation in vitro and in cells. Using microfluidics-based scanning to measure phase boundaries and microfluidic resistive pulse sensing (MRPS) to quantify size distributions of so-called pre-percolation clusters in subsaturated solutions [60], [61], we characterized the differential effects of molecular grammars on the assembly of synapsin-1 condensates. Finally, using pH-sensitive fluorescent dyes, we demonstrate that protons partition preferentially into dense phases of synapsin-1 condensates, thus generating a distinct chemical environment. Overall, our data point to the existence of conserved, sequence-specific molecular grammars in synapsin condensates, which could be crucial to ensuring the functional co-existence of multiple synaptic condensates.

## Results

### IDRs of SV proteins are defined by conserved and complementary molecular grammars

Previous studies have shown that IDRs of proteins that make up condensates are defined by condensate-specific molecular grammars [15], [18] [19] [21] [22] [23] [48] [62]. Here, molecular grammars refer to the evolutionarily conserved, non-random compositional biases and binary sequence patterns within an IDR sequence [19] [23] [44] [55] [63]. We find that approximately 50% of resident SV proteins [64] and SV-accessory proteins [65] contain IDRs. To assess if distinct molecular grammars exist within the IDRs of resident SV proteins and soluble SV-accessory proteins, we deployed NARDINI+, which calculates the z-scores of 90 sequence features for each SV protein [19] [23] [44] [55]. Here, both compositional and patterning features were examined. The z-scores quantifying compositional biases were calculated by referencing against the human IDRome, whereas the z-scores for binary sequence patterns were calculated by referencing against 10^5^ random sequences of the same composition.

We find that IDR grammars of resident SV proteins and accessory proteins are distinct from other condensates and from one another (**Figure 1A**). IDRs of resident SV proteins are enriched in negatively charged and aromatic residues when compared to the human IDRome (**Figure 1A**). These include the cytoplasmic tails of SYP and synaptogyrin-1 (SYNGR1) which are enriched in aromatic residues, and the cytoplasmic tails of the vesicular glutamate transporters (VGLUTs) and the synaptic vesicle glycoproteins (SV2s) which are enriched in both negatively charged and aromatic residues (**Figure 1B**). In contrast, soluble SV-accessory proteins are enriched in prolines and blocks of polar and proline residues that are segregated from one another (**Figure 1A**). Additionally, many of these proteins are positively charged with a preference for arginine over lysine residues, and this includes the synapsins (SYNs) (**Figure 1B**).

**Figure 1:**
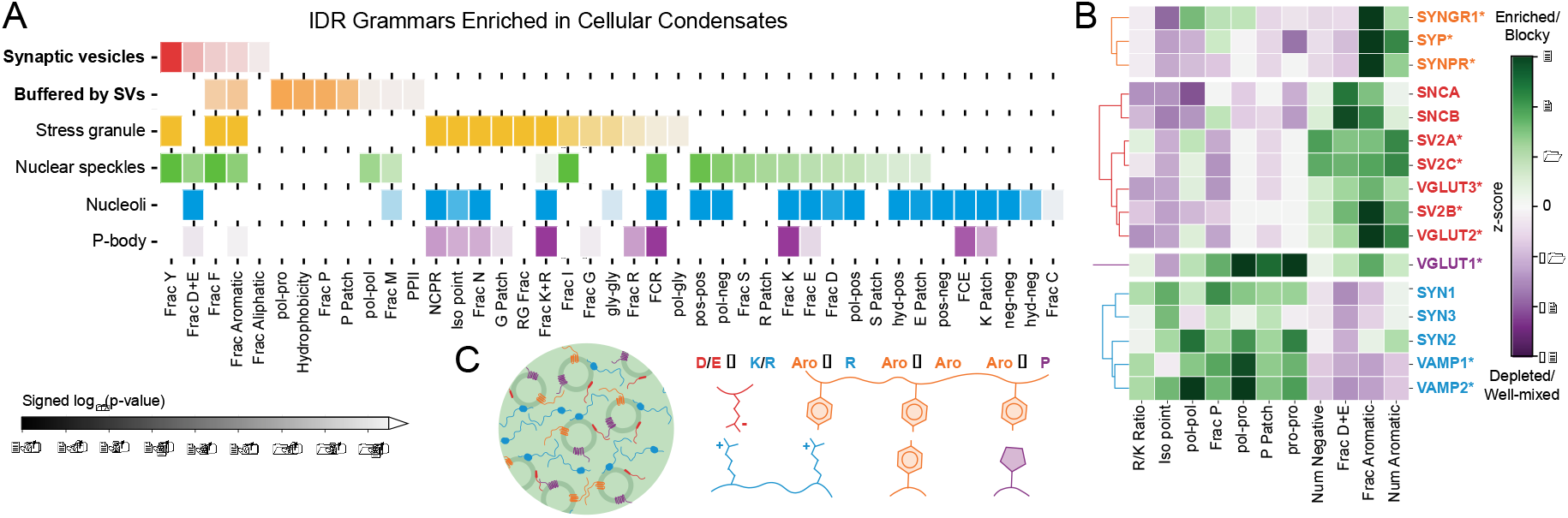
IDRs of proteins that make up synaptic condensates feature distinct molecular grammars. **A**. NARDINI+ analysis shows that IDR grammars of SV proteins are distinct from those of IDRs in other condensates. P-values were calculated using the two-sample Kolmogorov-Smirnov test between the distribution of z-scores for IDRs in a given condensate versus the distribution of z-scores for all remaining IDRs of length ≥30 in the human proteome **B**. NARDINI+ analysis of main SV proteins. Compositions and sequence patterns of IDRs of pre-synaptic proteins fall into distinct clusters denoted by dendrograms and different colors of the protein names. Here, * denotes SV resident proteins. **C**. Schematic illustrating the chemical complementarity across IDRs within proteins that are a part of SV clusters.

The molecular grammars of IDRs in resident transmembrane and accessory SV proteins are conserved across metazoans from lamprey, the oldest vertebrate predecessor, to humans (*SI Appendix* **Figure S1**). The distinct and evolutionarily conserved IDR grammars enriched in the resident and accessory SV proteins suggest that complementary interactions among IDRs may mediate the formation and regulation of SV condensates. Specifically, complementary charge interactions, interactions between aromatic residues, proline and aromatic interactions, and cation-pi systems, especially arginine and aromatic residues may mediate both homotypic and heterotypic interactions (**Figure 1C**) [8] [66].

### In vitro and in cell assessments of the effects of conserved molecular grammars

To assess the impact of distinct molecular grammars on SV condensation, we focused on synapsin-1, a key protein that drives the assembly of SV condensates [1]. Synapsin-1 has an N-terminal IDR, a folded substrate binding domain, and a C-terminal IDR. The C-terminal IDR is enriched in segregated blocks of polar / proline residues and shows a clear preference for arginine over lysine (**Figure 1B**). To test the impacts of these distinctive grammars, we designed two variants of synapsin-1 where the designs focus on the C-terminal IDR. In the SCR variant, we scrambled the sequence of the IDR to disrupt proline / polar blocks while keeping the remaining sequence and the overall amino acid composition fixed (**Figure 2A**). In the RtoK variant, arginine residues within the IDR were substituted with lysine (**Figure 2B**). This mutant was designed to alter the chemical specificity while maintaining the basic nature of the residues and their patterning within the sequence.

**Figure 2:**
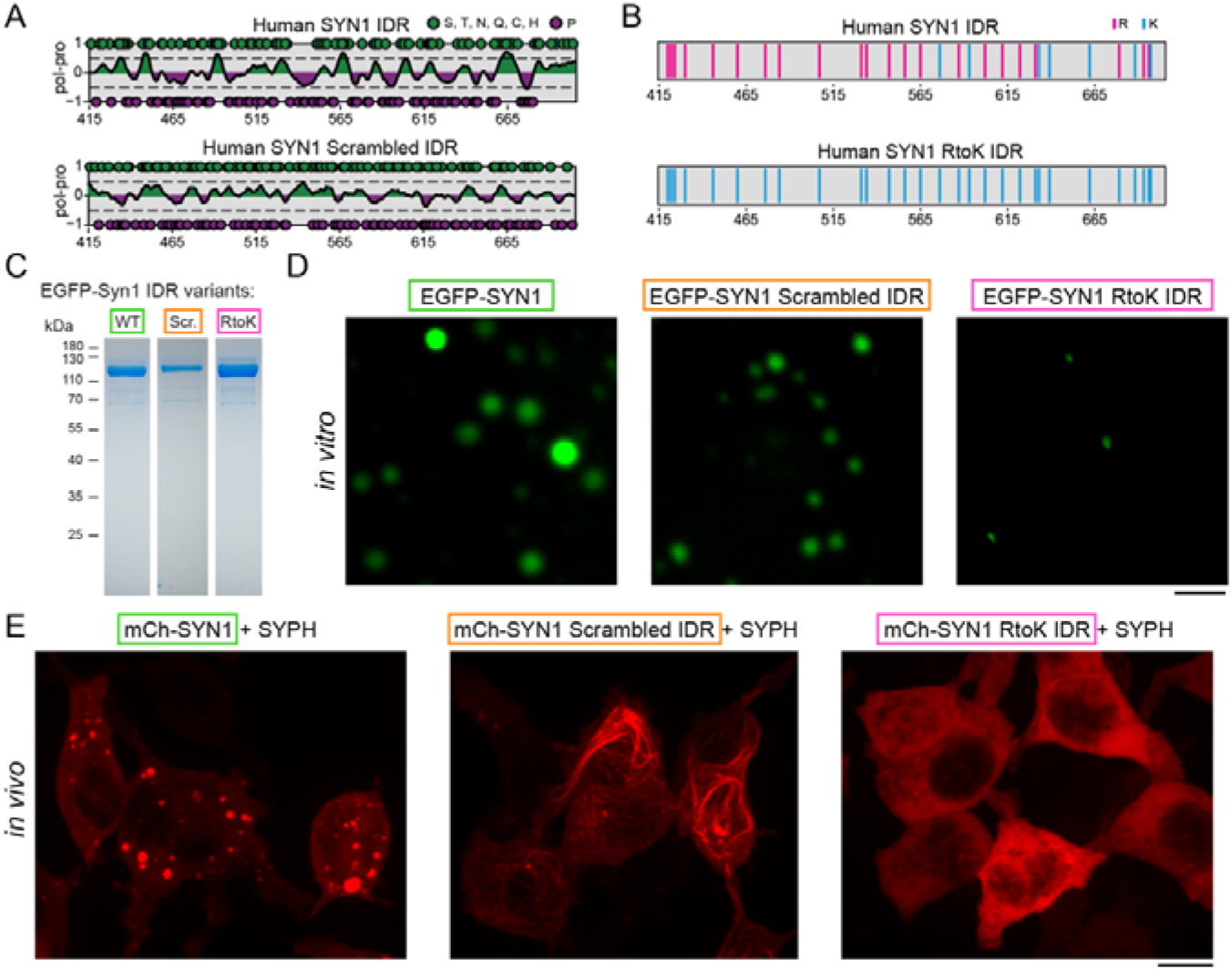
Disruption of the conserved molecular grammars within the C-terminal IDR of synapsin-1 can weaken or abrogate condensate formation in vitro and in cells. **A**. Schematic showing the synapsin-1 scrambled IDR in which polar and proline blocks are evenly distributed to reduce blockiness. **B**. Schematic of the synapsin-1 IDR showing the substitution of Arg with Lys. **C**. Coomassie gel indicating the final recombinant proteins used for the biophysical assays. **D**. Microscopy images of EGFP-synapsin-1 (8 μM) with WT, SCR, and RtoK IDR reconstituted in 25 mM Tris-HCl (pH 7.4), 0.5 mM TCEP, 150 mM NaCl and 3% PEG 8,000. Scale bar, 5 μm. **E**. Images of HEK cells co-expressing mCherry-tagged synapsin-1 WT (left), scrambled IDR (middle), or RtoK IDR (right) with untagged synaptophysin. Mutants disrupted the formation of normal synapsin condensates. IDR scramble formed filaments resembling cytoskeleton, and RtoK completely abolished condensation. Scale bar: 10 μm.

For in vitro characterizations of phase behaviors, we cloned, expressed, and purified the WT and the two mutants. All proteins have an EGFP fluorescent tag (**Figure 2C**, *SI Appendix* **Figure S2**). In the presence of 3% PEG 8000 we used microscopy to characterize the micron-scale assembles formed by 8 μM of each of the three proteins in 25 mM Tris-HCl (pH 7.4), 0.5 mM TCEP, and 150 mM NaCl. The WT and the two variants showed stark differences in their ability to form condensates. WT synapsin-1 readily formed condensates of 2-3 μm in diameter, whereas the SCR construct formed smaller condensates of up to ∼1 μm in diameter. These micron-scale condensates were not observed for the RtoK mutant construct under identical solution conditions (**Figure 2D**). It is clear that substitution of arginine to lysine weakens homotypic interactions, due to altered hydrophobicity and weakening of pi-pi interactions, and abolishes the formation of micron-scale assemblies [13] [15] [17] [24] [67] [68] [69] [70] [71] [72].

SYP is an integral SV membrane protein that is responsible for the generation of small vesicles. SYP contains a cytoplasmic IDR enriched in aromatic residues (**Figure 1B**) [73]. We conjectured that the arginine-, proline-, and polar-rich molecular grammar of synapsin-1 IDR and the aromatic-rich molecular grammar of SYP IDR should enable complementary cation-pi-pi interactions [66]. We tested this hypothesis by co-expressing the synapsin-1 variants with SYP in cells. While co-expression of mCherry-tagged WT synapsin-1 with SYP is known to recapitulate co-condensation in cells [6], condensates failed to form when we co-expressed the SCR variant of synapsin-1. Instead, this variant showed a gain-of-function phenotype with an apparent ability to track along actin filaments (**Figure 2E**).

The substitution of arginine with lysine in the synapsin-1 IDR also abrogated co-condensation with SYP in cells (**Figure 2E**). Therefore, even though lysine and arginine are basic residues, co-condensation with SYP requires the pi-like character of arginine. Therefore, arginine and lysine are not interoperable in the context of the C-terminal IDR of synapsin-1. Furthermore, the interactions between SYP and synapsin-1 appear to be dominated by complementary cation-pi-pi interactions rather than complementary electrostatic interactions alone [8]. Overall, the loss of the pi-like character of arginine residues compromises phase separation of synapsin-1 in vitro and co-condensation with SYP in cells.

### Synapsin-1 IDR grammars directly affect driving forces for phase separation

For rigorous quantification of driving forces for phase separation, we mapped comparative phase boundaries for the synapsin-1 variants using a microfluidics-based “PhaseScan” approach [74] (**Figure 3A**, *SI Appendix*). Phase separation is defined by the presence of threshold concentrations or threshold solubility products depending on whether homotypic versus heterotypic interactions drive the process [75], [76]. Measuring phase boundaries requires a scan in concentration space to delineate the one-phase and two-phase regimes. Synapsin-1 WT, synapsin-1 SCR, and synapsin-1 RtoK each showed distinct phase boundaries (**Figure 3B**). Scrambling the C-terminal IDR of synapsin-1 and/or R-to-K substitutions within the IDR significantly weakened the driving forces for phase separation. This is clear from a rightward shift of the phase boundaries towards higher PEG concentrations, a decrease in the slope, and changes in the shape of the coexistence curves (**Figure 3C**). The altered shapes of phase boundaries and weakened sensitivity to changes in protein and PEG concentrations point to decreased dominance of homotypic interactions [77] engendered by scrambling the sequence and / or arginine to lysine substitutions.

**Figure 3.**
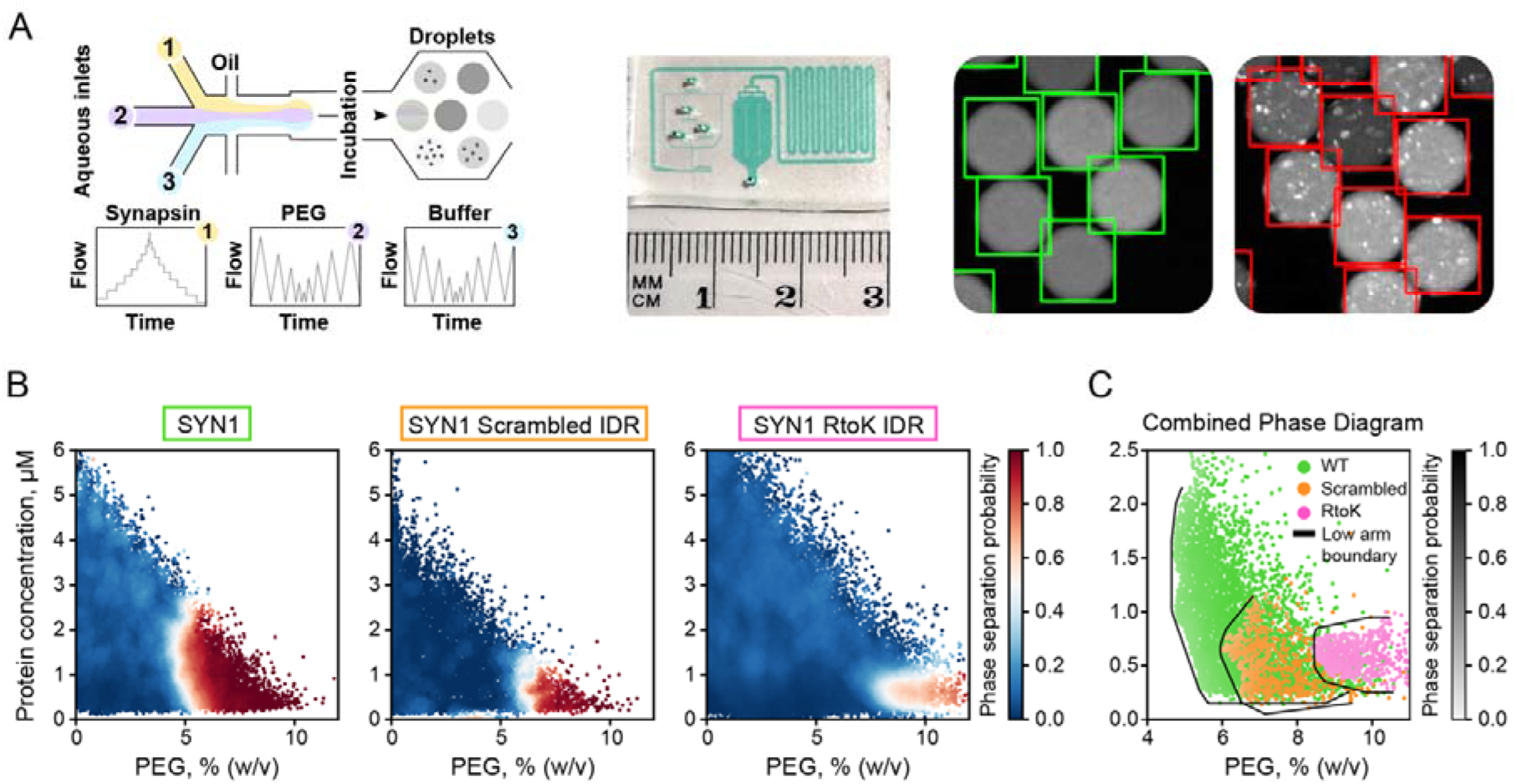
Molecular grammars of synapsin-1 IDR influence synapsin-1 condensation. **A**. Left: scheme of the microfluidic chip. The insets 1, 2, and 3 are flow rates of synapsin-1, PEG and buffer. Middle: an exemplary photograph of the chip (filled out with a colored solution to highlight the device features). Right: micrographs of individual droplets containing either mixed (green squares) or phase separated (red squares) solution. Classification is performed on each individual droplet as described in Methods. **B**. Low concentration arms of phase boundaries of EGFP-synapsin-1 WT (left, N = 29,187 points), EGFP-synapsin-1 scrambled IDR (middle, N = 13,203 data points), or EGFP-synapsin-1 RtoK IDR (right, N = 64,675 data points). Data points correspond to individual microdroplets classified as phase separated (red) or homogeneous (blue). The heatmap quantifies the phase separation probability. **C**. Combined phase boundaries from **A** showing the low concentration arm. The boundaries are drawn using an algorithm, that segregates the measured points that belong to the one-phase versus two-phase regimes.

### Impact of synapsin-1 IDR grammars on pre-percolation clusters in subsaturated solutions

Condensation combines reversible binding, phase separation and percolation [15] [23] [59] [78] [79]. A signature of percolation is the formation of heterogeneous distributions of pre-percolation clusters in subsaturated solutions [60] [61] [80]. The latter refers to concentrations that are below the threshold for phase separation. Pre-percolation clusters form as the result of heterogeneities of so-called sticker versus spacer mediated interactions [60] [61]. Stickers refer to cohesive domains, motifs, or residues and spacers are interspersed between stickers [59]. The average sizes of pre-percolation clusters typically increase with increasing concentrations. Furthermore, the distributions of cluster sizes are heavy-tailed implying that the dominant clusters are oligomers, whereas the lower likelihood species are clusters comprising hundreds of molecules [60], [61].

Percolation and phase separation are governed by two different sets of interactions [59] [78] [81] [82]. The strengths of reversible associations between stickers determine the percolation threshold and the concentration-dependent size distributions of pre-percolation clusters. These clusters grow continuously with concentration and become system-spanning above the percolation threshold [59] [79]. In contrast, phase separation is governed by solubility considerations, which are quantified by the Flory-Huggins χ parameter [83] [84]. A requirement for phase separation is that χ > 0.5, and the solubility limit, which defines the saturation concentration or solubility product is set by the magnitude of χ.

Recently, Kar et al., showed that phase separation is realized in aqueous solvents when the sizes of pre-percolation clusters cross a threshold size of 800 nm – 1 μm [61]. Overall, the valence of stickers and the strengths of attractive interactions between stickers set the percolation threshold and the concentration-dependent evolution of the size distributions of pre-percolation clusters. The Flory-Huggins χ parameter, which is governed jointly by the solvation preferences of spacers that are interspersed between stickers and sticker-sticker interactions will determine the driving forces for phase separation. The overall implication is that the contributions to phase separation and percolation can be (de)coupled [82], and this is evident in comparing the phase behaviors of the WT and designed variants of synapsin-1.

We deployed microfluidics resistive pulse sensing (MRPS) [85] [86] to quantify the size distributions of clusters in subsaturated solutions [60]. In MRPS, individual particles in a sample flow through a nano-constriction across a potential drop in a microfluidic cartridge. The changes to the electrical resistance over the transit time of the particle are used to calculate the particle concentration and diameter, respectively [86]. This label-free method enables the quantification of sizes ranging from 70 nm to 10 μm.

Substitution of arginine to lysine alters the size distributions of pre-percolation clusters showing a pronounced shift toward smaller species (**Figure 4A,4B**). This points to weakened self-associations that, in turn, weaken clustering in subsaturated solutions and the driving forces for phase separation. The latter manifests as increased threshold concentrations for phase separation, requiring higher concentrations of PEG (**Figure 3B-C**). In contrast, even though the driving forces for phase separation are weakened when the sequence of the IDR is scrambled in the SCR variant, clustering in subsaturated solutions is equivalent for synapsin-1 WT and synapsin-1 SCR. This implies that the driving forces for forming pre-percolation clusters and phase separation can be decoupled by the scrambling of proline and polar blocks While percolation appears to be drive purely by the valence i.e., the number of proline and polar residues, the solubility limit is altered by scrambling these residues across the entirety of the sequence. This observation dovetails with reports of how polar tracts lower the solubility limits of IDRs [87] [88].

**Figure 4.**
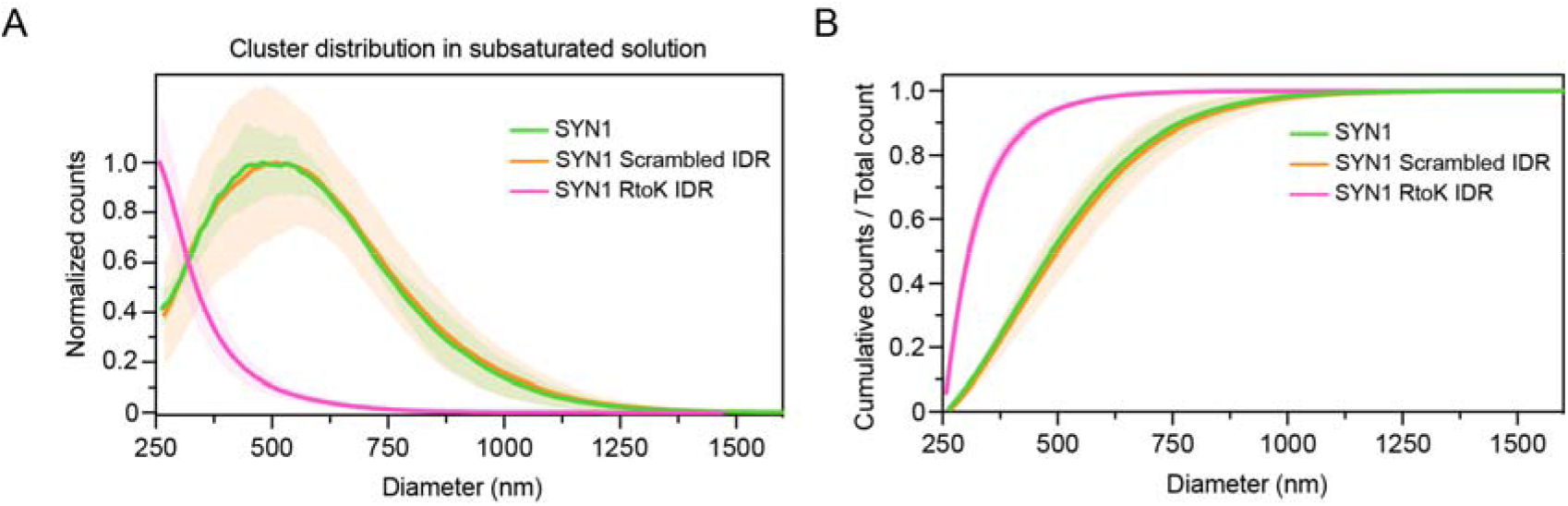
Comparative assessments of cluster size distributions in subsaturated solutions. **A**. Cluster size distributions in subsaturated solutions for synapsin-1 WT (green), SCR (orange), and RtoK (magenta) variants. Measurements were performed at equivalent conditions (8 μM for each synapsin-1 variant with 3% PEG 8000 in 25 mM Tris-HCl (pH 7.4), 150 mM NaCl, 0.5 mM TCEP) using a C-2000 cartridge in an nCS1 instrument. The plot shows the counts that are normalized to the maximum value as a function of the particle diameter. **B**. Cumulative counts for cluster size distributions shown in A. Error bars (shaded) represent standard errors of the mean values.

### Interphase pH gradients in synapsin-1 condensates

Biomolecular condensates are defined by two or more coexisting phases. Chemical potentials of all solution components are equalized across the phase boundary, and preferential interactions of solutes or ions such as protons with molecules in the dense or dilute phase can generate passive interphase potentials [89] and pH gradients [23]. We asked if phase separation of synapsin-1 generates a pH gradient. This is because maintaining a pH gradient is likely to be essential for vesicular ATPase functions and the loading of neurotransmitters into SVs [90] [91].

To determine if there is an interphase pH gradient and assess how this might be affected by designed mutations to the IDR of synapsin-1, we employed ratiometric pH-sensitive dyes [23], Seminaphtharhodafluor (SNARF) dyes, specifically SNARF-1. If protons preferentially bind to molecules in the dense phase, then at equilibrium, there will be a passive, interphase proton (pH) gradient, making the dense phase more acidic than the coexisting dilute phase. The converse will be true if protons bind preferentially to molecules in the dilute phase.

Using the ratiometric pH-sensitive dye SNARF-1 (**Figure 5A-5C**), we found that synapsin-1 WT, SCR, and RtoK showed equivalent abilities to accumulate protons in dense phases (**Figure 5D**). Therefore, while molecular grammars in synapsin-1 control driving forces for condensation, the interphase pH gradients remain unchanged, presumably because sequences have an identical net charge [23].

**Figure 5.**
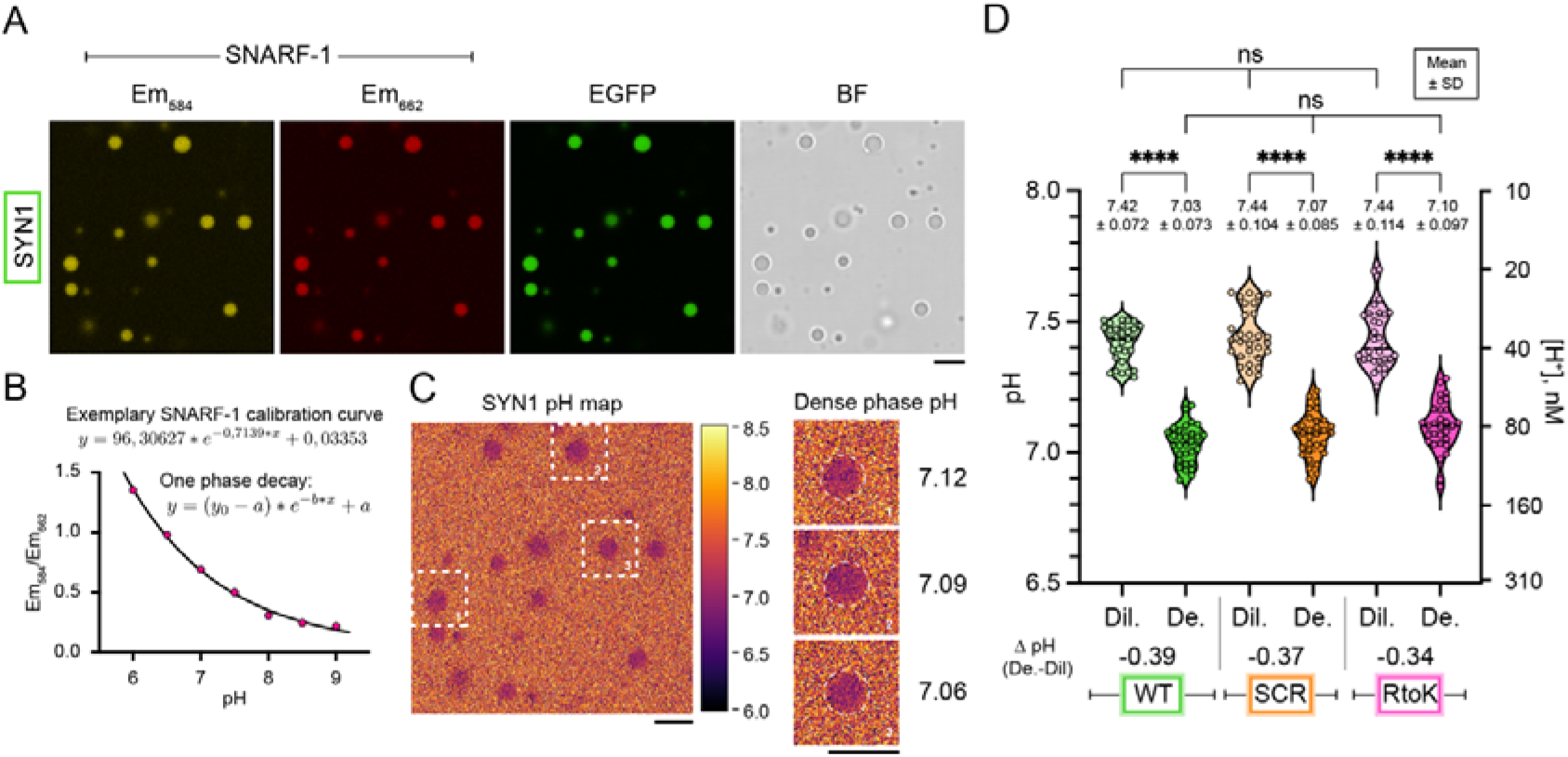
Synapsin-1 condensates sequester protons and lower pH. **A**. Representative micrographs of EGFP-synapsin-1 condensates (8 μM protein in 3% PEG 8,000) co-incubated with the ratiometric pH-sensitive reporter SNARF-1. SNARF-1 was excited at 561 nm and recorded at 582-586 (yellow) and 660-664 nm (red). Scale bar, 5 μm. **B**. Carboxy SNARF-1 calibration curve for reconstitution buffer (25 mM Tris-HCl, 150 mM NaCl, 0.5 mM TCEP) at different pH (pH 6 – pH 9) on a single plane. Data were fit to a one-phase-decay function. **C**. Left: Example micrograph indicating a pH map for EGFP-synapsin-1 WT condensates calculated by applying the z-slice calibration curve. Heatmap represents pH. Right: Magnified regions of selected condensates (dashed line) with indicated average pH values of the dense phase. Scale bar, 5 μm. **D**. pH analysis for the dense and dilute phases of condensates for synapsin-1 WT (green), SCR (orange), and RtoK (magenta) variants. Average pH values are shown in the graph with standard deviation. Violin plots with median and quartiles indicated by the dashed lines. (One-way ANOVA, ****: p<0,0001).

## Discussion

Our study shows that the conserved molecular grammars within the C-terminal IDR of synapsin-1 influence condensation via homotypic interactions in vitro and co-condensation with SYP in cells. Specifically, arginine residues are necessary to drive homotypic and heterotypic interactions that cannot be accounted for purely due to arginine being a basic residue. Instead, arginine is a stronger sticker than lysine due to its pi-like character and stronger cation-pi-pi interactions [66] that can engage with aromatic residues in synapsin-1 and other pre-synaptic proteins.

The blocky patterning of polar and proline residues is also necessary for homotypic associations and for proper cellular interactions of synapsin-1. Blocks of polar residues may contribute to homotypic interactions [19] [87] [88], whereas proline-rich regions can contain motifs that interact with SH3 domain-containing pre-synaptic proteins, such as intersectin [1] [92]. In contrast, the formation of pre-percolated clusters does not depend on the segregation of polar and proline residues.

Our work demonstrates how insights from computational analysis, uncovering evolutionarily conserved molecular grammars, can be tested experimentally via designed mutations [20][21][22]. This approach bypasses the often-used approach of deleting IDRs to assess their contributions – an approach that rests on the notion that IDRs engage mainly in non-specific interactions [10] [11]. Instead, we show how systematic designs that modulate conserved grammars can be brought to bear to investigate how these regions contribute to phase behavior.

Although the focus of our study has been on the IDR of synapsin-1, our approach can be applied to study how interactions between various pre-synaptic proteins modulate the formation and regulation of SV condensates. Approximately half of the SV-resident and soluble SV accessory proteins contain IDRs, and these two sets of pre-synaptic proteins contain distinct and complementary IDR grammars (**Figure 1A**). We propose that SV clustering is regulated by interactions involving complementary molecular features that are distributed across IDRs of different pre-synaptic proteins. Previous studies support this hypothesis for heterotypic buffering as the SV resident proteins SYP, VGLUT1, and VAMP2 and the accessory protein α-synuclein have all been shown to modulate SV clustering [93] [94] [95] [96] [97] [98]. Additionally, a recent study showed that resident proteins SYP, SYNGR1, and SYNPR can all co-condense SVs with synapsin-1 [99]. Our analysis shows that the cytoplasmic C-terminal IDRs of these three proteins cluster separately from the other SV condensate-modulating pre-synaptic proteins due to their enrichment in aromatics but lack of other grammar features (**Figure 1B**). In contrast, VAMP2, which houses an IDR with a similar molecular grammar to synapsin-1 cannot form condensates with synapsin-1 in cells [6]. However, VAMP2 can be recruited into co-condensates of synapsin-1 and SYP. These results are consistent with complementary grammars, including cation-π interactions, driving SV condensation, whereas like grammars may play more modulatory roles.

In the context of the pre-synapse, the results presented here and investigations of the synergies of molecular grammars in synaptic boutons [5] are likely to enable new insights into our understanding of the neuronal transmission that is relevant in healthy brains and diseases characterized by the loss of functional synapses.

## Material and Methods

### Analysis of IDRs using NARDINI+

Sequence grammar analysis was performed using NARDINI+ as described in [23]. Briefly, NARDINI+ calculates z-scores of compositional and patterning grammar features previously shown to be important for IDR conformation, interactions, and function [48]. Five additional compositional grammar features were examined that were not described in King et al. These include the number of positive, negative, polar, aliphatic, and aromatic residues, giving a readout on valence compared to the full IDRome. For a given IDR, compositional z-scores are calculated by extracting the compositional feature of the given IDR and using the mean and standard deviations from the full human IDRome (24508 IDRs). The human IDRome contains all IDRs of length ≥ 30, extracted from the UniProt human proteome (UP000005640) and MobiDB predictions [100] [101]. To calculate the patterning z-scores for a given IDR, NARDINI+ is performed [55]. Here, the asymmetry of patterning of two residue types in a given IDR sequence is compared to the mean and standard deviations of 10^5^ random sequences of the same composition.

### Extraction of condensate-specific enriched grammar features

Condensate-specific protein lists were extracted from various sources and cross-referenced with the human IDRome to extract condensate-specific IDRs. As described in [23], proteins localized to nucleoli and nuclear speckles were extracted from the Human Protein Atlas (HPA). Proteins localized to P-bodies and stress granules were extracted from [102] and [103], respectively. Synaptic vesicle proteins were extracted from [64] using the list of “SV-resident” proteins leading to 125 IDR sequences. Buffered by SV proteins were extracted from [65] (here paralogs were included) to yield 47 IDRs. To determine which grammar features were enriched in specific condensates, the human IDRome was split into two sets: (1) the current condensate-specific IDR set and (2) the remaining human IDRome. Then, the distributions for each of the 90 different z-scores were compared using the two-sample Kolmogorov-Smirnov test and a p-value was calculated. The signed log_10_(p-value) was calculated for p-values < 0.05. Specifically, −log_10_(p-value) was calculated if the condensate-specific mean z-score value was greater than the rest of the IDRome mean z-score value, and log_10_(p-value) was calculated if the condensate-specific mean z-score was smaller. Thus, positive values imply the given feature is enriched or blockier in the condensate-specific IDR set.

### Extraction of IDR grammar clusters within SV proteins

Grammar feature z-scores were calculated using NARDINI+ as described above for IDRs of SV proteins: SNCA, SNCB, SNCG, SYN1, SYN2, SYN3, SYP, SYNPR, SYNGR1, SV2A, SV2B, SV2C, STX7, VAMP1, VAMP2, VGLUT1, VGLUT2, and VGLUT3. Figure 1B shows a filtered set of enriched features for the four clusters having at least 3 IDRs determined by hierarchical clustering using the Euclidean distance and Ward’s linkage method [104].

### Evolutionary analysis of synapsin IDRs

The list of eukaryotic SYN1 orthologs was extracted from EggNOG [105]. The proteomes of 12 additional species were examined for orthologs if not included in the original EggNOG list. These include *Aptenodytes forsteri* (taxid: 9233), *Doryteuthis pealeii* (taxid: 1051067), *Lampetra fluviatilis* (taxid: 7748), *Leptonychotes weddellii* (taxid: 9713), *Marmota marmota marmota* (taxid: 9994), *Microcebus murinus* (taxid: 30608), *Octopus vulgaris* (taxid: 6645), *Petromyzon marinus* (taxid: 7757), *Physeter macrocephalus* (taxid: 9755), *Urocitellus parryii* (taxid: 9999), *Ursus americanus* (taxid: 9643), and *Ziphius cavirostris* (taxid: 9760) whose proteomes were extracted from UniProt. BLASTp was used to extract only the most similar ortholog per species compared to the full SYN1 sequence [100] [106]. Then, MUSCLE sequence alignment was utilized to align orthologous sequences [107]. From the alignments, Jalview was used to extract the IDR regions in orthologs corresponding to residues 415-705 of human SYN1 [108]. Only orthologous sequences of length ≥ 15 and housing no unnatural amino acids were kept. NARDINI+ was then run on all orthologous sequences where the human IDRome was used as the prior to extract the z-scores of the compositional features. Overall, 152 species were analyzed.

### Calculation of mean net residue type profiles

Here, we consider residue types pol≡{C, H, N, Q, S, T} and pro≡P. For a given sequence, the fraction of pol residues minus the fraction of pro residues is calculated for each sliding window of length five. Then, the values from all sliding windows that contain a given residue are averaged to yield a residue specific mean net residue type value. These values are plotted in Figure 2A and S1C.

### Cloning of synapsin-1 IDR variants

The human synapsin-1 coding sequence is based on the NM_006950.3 NCBI reference Sequence. The complexity of the sequence, such as GC-rich regions, was manually altered to allow efficient gene synthesis while leaving the protein sequence unchanged (NP_008881.2). An endogenous SacI restriction site at codons E408-E409 (GAGCTC) was silenced by altering the nucleotide sequence to GAGCTG. To allow for easy subsequent exchange of the synapsin-1 IDR, a BamHI restricition site was added at codons G389-S390 by changing the nucleotide sequence from GGTTCC to GGATCC. The optimized human synapsin-1 codon sequence and the variant IDR sequences (a.a. 389-705) were created by gene synthesis (Eurofins, HIFI DNA assembly). Subsequently, IDR fragments starting from a.a. 389 were subcloned using the restriction sites BamHI and SacI, yielding sequences of full-length Synapsin 1 containing IDR variants. The resulting full-length synapsin-1 sequences (WT, scrambled IDR, RtoK IDR) were either subcloned into a pCMV expression plasmid as a C-terminal fusion protein to mCherry for microscopy [4] by Kpn2I and SacI or subcloned into a His14-SUMO_Eu1-EGFP-MCS plasmid for protein purification by BglII and KpnI [3]. All constructs were verified by Sanger sequencing.

### Cell culture and transfection

HEK 293T cells were grown in Dulbecco’s Modified Eagle’s Medium GlutaMAX Supplement (10566016, Gibco) supplemented with 10% fetal bovine serum (12106C, Sigma), 1% MEM Non-essential Amino Acid Solution (M7145, Sigma) and 1% penicillin/streptomycin (15140122, Gibco) at 37 °C and 5% CO_2_. For imaging, cells were seeded on 25 mm cover glasses. Cells were transfected with a total of 2 μg of plasmid DNA using Lipofectamine 2000 (11668500, Invitrogen) following the manufacturer’s instructions. In brief, the final transfection mixture contained 3 μL of lipofectamine with 2 μg of total plasmid DNA in 200 μL OptiMEM (31985070, Gibco). The transfection mix was incubated for 20 min at room temperature before adding to seeded HEK cells (confluency ∼ 60-70%) and incubated overnight at 37 °C and 5% CO_2_. For protein purification experiments, Expi293F™ suspension cultures (A14527, Gibco) were maintained in Expi293™ Expression Medium (A1435102, Gibco) according to the manufacturer’s protocol (37 °C, 8% CO2, 125 rpm). Transfection of suspension cultures (30 mL) was performed with 30 μg of plasmid DNA following the ExpiFectamine™ 293 Transfection Kit guidelines.

### Protein expression and purification of synapsin-1 IDR variants

EGFP-tagged synapsin-1 IDR variants were expressed in EXPI cells (HEK derivative) and purified by a two-step purification strategy consisting of NiNTA-affinity and size-exclusion chromatography followed by His-SUMO-tag removal, as described [3] [109]. Briefly, His14-SUMO_Eu1-synapsin-1 IDR variants were expressed in Expi293F™cells (Thermo Fisher Scientific) for three days following enhancement. Cells were lysed by three cycles of freezing and thawing in a buffer that contained 25 mM Tris-HCl (pH 7.4), 300 mM NaCl, 0.5 mM TCEP (buffer A) supplemented with EDTA-free Roche Complete protease inhibitors, 15 mM imidazole, 10 μg/mL DNaseI and 1 mM MgCl_2_. The purification steps were carried out at 4°C. Lysate clearance was done by centrifugation for 40 min at 30,000xg. The soluble supernatant was incubated with complete His-Tag Purification resin (Roche) in a polyprep column (Biorad) on a rotating platform for 1 h. Washing steps were done in buffer A with 15 mM imidazole, and elution was performed in buffer A with 400 mM imidazole. Fractions were concentrated using a 30 K MWCO protein concentrator (Millipore) and applied to size exclusion chromatography (Superdex™200 Increase 10/300, GE Healthcare) in a buffer containing 25 mM Tris-HCl (pH 7.4), 150 mM NaCl, 0.5 mM TCEP. Fractions that contained the His14-SUMO_Eu1-Synapsin 1 IDR variant were combined and subjected to tag cleavage with SENP_EuB SUMO protease digest overnight (protease:protein ratio of 1:50) at 4°C. After incubation, the reaction was supplemented with imidazole and NaCl (final concentrations: 15 mM and 300 mM, respectively). The cleaved His-SUMO-tag and His-tagged SENP protease were removed by passing the solution three times over preequilibrated (buffer A with 15 mM imidazole) complete His-Tag Purification resin. Proteins in the flow-through were subjected to buffer exchange using a PD-10 column (final buffer: 25 mM Tris-HCl (pH 7.4), 150 mM NaCl, 0.5 mM TCEP) and concentrated with a 30 K MWCO protein concentrator (Millipore). Proteins were snap-frozen in liquid nitrogen and stored at −80 °C until use.

### Microscopy and image analysis

For *in vitro* reconstitution of EGFP-Synapsin 1 IDR variants, 8 μM protein was coincubated with polyethylene glycol (PEG 8,000) at a final concentration of 3% in a buffer containing 25 mM Tris-HCl (pH 7.4), 150 mM NaCl and 0.5 mM TCEP. After the addition of PEG, the condensation reaction was transferred to a glass bottom dish (Cellvis D35-20-1.5-N). Imaging was performed on a Nikon spinning disk confocal CSU-X microscope, equipped with a Plan Apo λ 60x Oil objective and a pco.edge camera. The excitation wavelength was 488 nm for EGFP, and image analysis was performed using ImageJ2 (Version: 2.9.0/1.53t). For live-cell confocal imaging in HEK cells an Andor DU-888 X-9798 camera and an excitation wavelength at 561 nm were used for mCherry-tagged synapsin-1 IDR variants.

### The PhaseScan approach for mapping phase diagrams

The microfluidic devices were fabricated using standard soft-lithography [74] [110]. Devices were designed using the AutoCAD (AutoDesk) software and then printed on a photomask (Micro Lithography). To obtain a master with microchannels of approximately 50 μm height, SU-8 3050 (Kayaku Advances Materials) was poured onto a silicon wafer (MicroChem) and spun using a spincoater (Laurell Technologies) for 45 s at 3000 rpm. After soft-baking at 95 °C for 25 min, the wafer with the photomask placed on it was subjected to UV exposure for 60 s. Post-exposure baking at 95 °C for 5 min was then followed by developing with propylene glycol methyl ether acetate (PGMEA, Sigma-Aldrich) and a washing step with isopropanol (IPA, VWR Chemicals). To fabricate the devices, polydimethylsiloxane (PDMS, Sylgard 184) was poured over the master, degassed and then baked at 65 °C for 1.5 h. Subsequently, devices were cut out, and the corresponding holes for the inlets and outlets were punched. A plasma oven (Diener Electronic) was used to treat the devices and enable bonding them to glass slides. The last step was to apply hydrophobic surface modification to the devices using 2% (v/v) trichloro(1H,1H,2H,2H-perfluorooctyl)silane (Sigma-Aldrich) in hydrofluoroether HFE-7500 (Fluorochem).

The PhaseScan microfluidic platform was used to map high-density phase diagrams [74]. The following three aqueous solutions were prepared to be mixed on-chip: 6 μM stock of one of the three EGFP-labelled synapsin-1 variants in 25 mM Tris-HCl (pH 7.4), 150 mM NaCl, 0.5 mM TCEP; solution of the corresponding buffer and a solution of polyethylene glycol (PEG 8000) 10% (w/v) in the same buffer, barcoded with 2 μM Alexa Fluor 647 free dye. Additionally, an oil solution of HFE-7500 (Fluorochem) with 4% surfactant (RAN Biotechnologies) was used for water-in-oil droplet encapsulation. The flow rates of the three aqueous solutions were varied automatically using pressure-controlled pumps (LineUp Flow EZ, Fluigent) according to a pre-programmed flow profile such that each microfluidic droplet contained different concentrations of protein and PEG. The total flow rate of the aqueous solutions was kept constant at 60 μL/h, while the oil flow rate was 150 μL/h, generating droplets of approximately 70 μm in diameter. Droplets were incubated for 3 minutes before imaging in the same microfluidic chip at two wavelengths (488 nm and 647 nm) at the same time using an epifluorescence microscope (Cairn Research) equipped with a 10x air objective (Nikon CFI Plan Fluor) and a dichroic filter set (Cairn Research).

A semi-automated custom-written Python script (Python version 3.9.7) was used to analyze the images, i.e., to detect droplets and classify them as phase separated (1) or mixed (0) by employing a convolution neural network (CNN) [111]. To allow calculation of the concentration of the components in each droplet, images of calibration droplets containing only the stock solutions of protein or PEG were recorded. A double-gaussian fit was applied over the fluorescence intensity histograms of the calibration droplets to correlate the intensities back to the stock concentration [112]. Hence, each microfluidic droplet provides information for one data point on the phase diagrams presented as scattered plots. Phase separation probability was further determined for each data point by averaging over its neighboring points within a radius of 5% of the maximum limit of each axis. The boundaries were determined using a fitting based on support vector machine-based methods.

### Extracting low concentration arms of phase boundaries

First, we generated a two-dimensional histogram of all points with a phase separation probability of at least 0.55. To eliminate outliers, if there exists at least three data points for the scramble or RtoK sequences and five data points for the WT sequence in a given bin, then the x and y positions of that bin are kept. The boundary is then drawn using these x and y positions and calculating the convex hull with rounded corners.

### Measurements of cluster distribution in the subsaturated solutions

Microfluidic Resistive Pulse Sensing (MRPS) measurements were performed on an nCS1 instrument (Spectradyne LLC, Signal Hill, CA, USA). 8 μM of the purified synapsin construct was mixed with 3% PEG 8,000 in the reaction buffer (25 mM Tris-HCl (pH 7.4), 150 mM NaCl, 0.5 mM TCEP) in an Eppendorf tube to induce phase separation. 5 μL of samples were measured using C-2000 cartridges, with a measurement range of 250–2000 nm. At least three acquisitions (triplicates) were collected and combined for analysis, with each acquisition being collected over 10 min. Control experiments were performed with 3% PEG 8,000, where no significant abundance of particles was detected. The particle size distributions from the triplicate experiments were averaged and the resulting curve was smoothed using the LOWESS method in Prism 10 (GraphPad, Boston, MA, USA).

### Analysis of pH in synapsin-1 condensates

The ratiometric pH-sensitive dye, 5-(and-6)-Carboxy SNARF™-1 (C1270, Invitrogen), was dissolved to a final concentration of 10 mM in ddH_2_O. For calibration, 25 mM Tris-HCl, 150 mM NaCl, and 0.5 mM TCEP buffer were prepared at different pH values (pH 6.0-9.0, steps: 0.5 pH units). All calibration curves were taken on the same day as the samples with synapsin-1 condensate were analyzed (all in triplicates). In brief, the calibration buffer was mixed with 20 μM final SNARF-1 dye (5 μL final reaction volume) and placed on a glass bottom dish (Cellvis D35-20-1.5-N). Imaging was performed on a Nikon laser scanning confocal AX NSPARC microscope (Ti2, AX camera, Galvano scanner, 2x line averaging, 1024×1024 px resolution, 1.0 μs dwell time) equipped with a Plan Apo λD 60x oil OFN25 DIC N2 objective. Ratiometric imaging of the SNARF-1 dye was performed by excitation at 561 nm and simultaneous readout of the two GaAsP PTMs with different freely tunable emission bands. The first band was set to 582-586 nm (center wavelength: 584 nm), and the second band was set to 660-664 nm (center wavelength: 662 nm). A total of 20 z-stacks were acquired from the cover glass surface into the buffer solution with a step size of 0.20 μm (total measured relative z-height 4.0 μm). Over the entire pH range, the calibration curves were acquired for each z-stack at the same distance from the surface of the coverglass and fitted to a one-phase decay function using GraphPad Prism 10.

For the pH analysis of EGFP-synapsin-1 IDR variants, 8 μM protein was coincubated with 3% PEG 8,000 and 20 μM SNARF-1 dye in a buffer containing 25 mM Tris-HCl (pH 7.4), 150 mM NaCl and 0.5 mM TCEP. Ratiometric z-stacks were recorded from the cover glass surface with the same step size and ratiometric imaging settings as for the calibration curves. Transmission detection was acquired using the 561-nm laser line. Additionally, imaging of EGFP was performed by excitation at 488 nm and emission at 499-551 nm. Of note, while synapsin-1 WT and scrambled IDR condensates formed readily within minutes, synapsin-1 RtoK IDR variant condensate formation was forced by extending the incubation time on the cover glass to one hour.

For image analysis, each pixel of the 584-nm emission image was divided by the corresponding pixel of the 662-nm emission image for each individual z-stack. The resulting ratio z-stack series was converted to a pH stack by applying the corresponding z-slice calibration curve to each matching ratio z-slice. All measurements were performed in three independent reconstitutions with 10 condensates selected from each single reconstitution. A dense phase ROI was outlined with the segmented line tool in ImageJ, and a corresponding circular dilute phase ROI (diameter: 4.3 μm) was placed 2 μm away from the dense phase. ROIs were selected on a single z-plane. Mean pH values of both dense and dilute phases were plotted as violin plots. One-way ANOVA tests (multiple comparisons, Tukey, alpha=0.05) were performed for statistical analysis in GraphPad Prism 10 (GraphPad, Boston, MA, USA).

## Supplementary Figures

**Figure S1.**
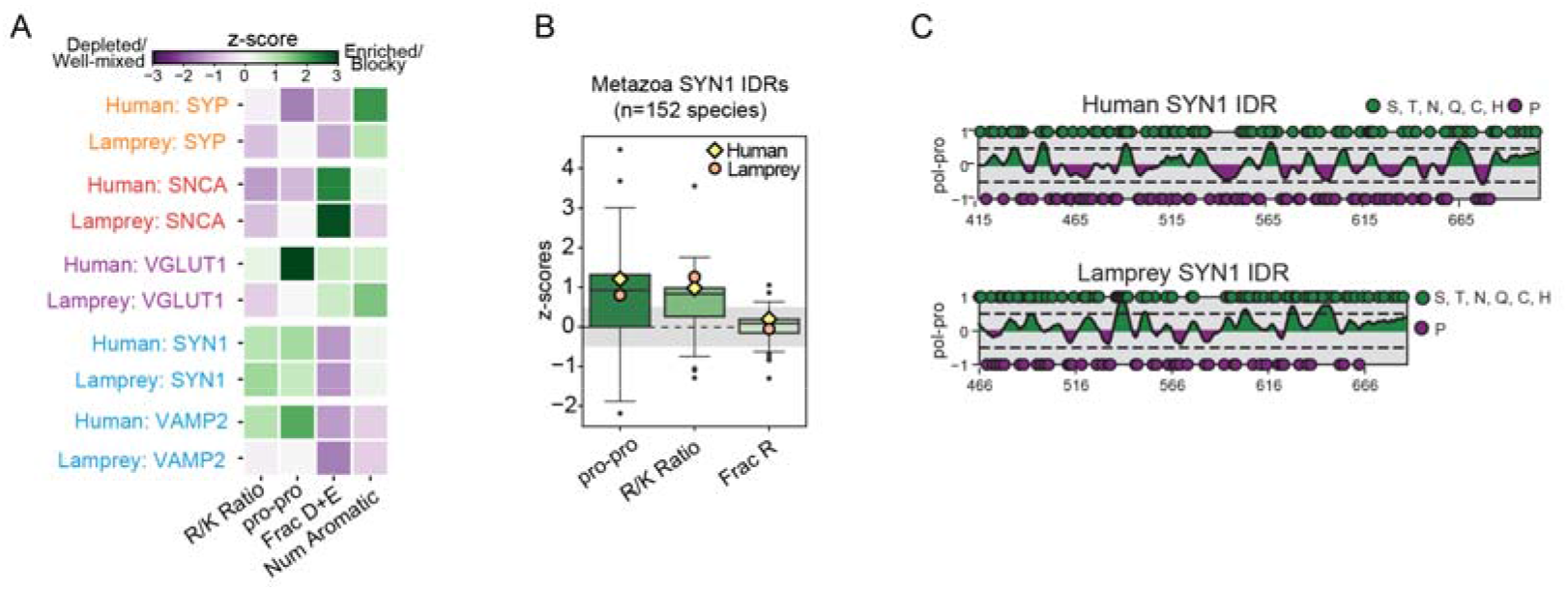
Chemical specificity within IDRs is highly conserved. **A**. IDRs of human and lamprey SV proteins have similar chemical features. **B**. Distribution of z-scores across 152 synapsin-1 orthologs. The molecular grammars are highly conserved across 152 metazoans including from lamprey, the oldest vertebrate predecessor, to humans. Synapsin-1 IDR grammar features, including proline blocks and Arg to Lys ratio are enriched, whereas Arg fraction is not enriched. **C**. Blockiness of polar and proline residues is conserved in the lamprey synapsin-1 IDR.

**Figure S2.**
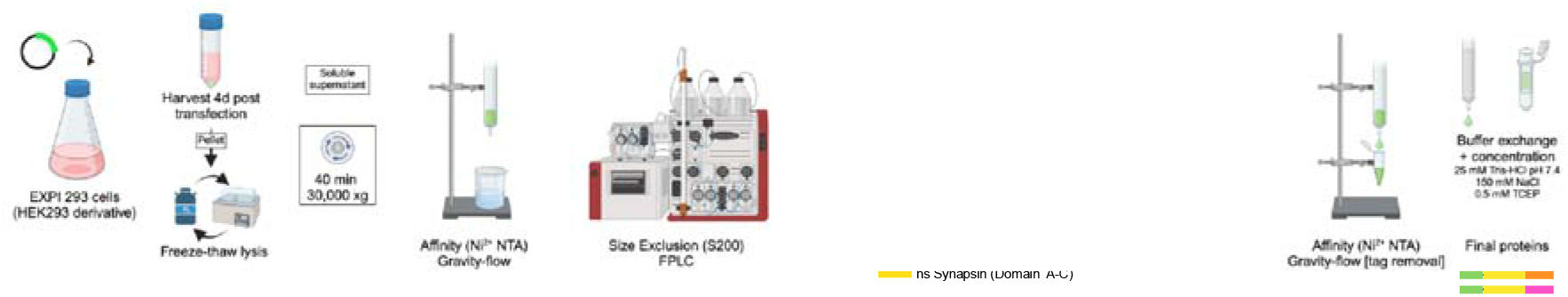
Scheme of the purification protocol for synapsin-1 IDR (WT) and the two variants SCR and RtoK. Scheme generated with BioRender and Adobe Illustrator.

## Acknowledgments

We thank the AMBIO Facility at Charité for microscopy. This work is supported by DZNE, grants from the DFG (MI 2104 and SFB1286/B10 to DM); ERC Grants (MemLessInterface 101078172 to DM, DiProPhys 101001615 to TPJK); the NIH (NIA 2RF1 NS078165-12 to JRM, NINDS R01NS121114 to RVP, and F32GM146418-01A1 to MRK); and AFOSR (FA9550-20-1-0241 to RVP). CH is supported by the German Dementia Association.

## Author Contributions

CH performed reconstitution experiments. KMR did the computational analyses. CH, IAE, HA performed microfluidic experiments and analyzed the data; CH and MKS performed MRPS measurements. CH, MRK, AP performed pH measurements. CH and JVT performed cell experiments. JRM and TPJK contributed to experimental designs. RVP and DM designed the study and wrote the paper.

## References

[1] D. Milovanovic, Y. Wu, X. Bian, P.D. Camilli, A liquid phase of synapsin and lipid vesicles, Science 361 (2018) 604 607. 10.1126/science.aat5671.

[2] A. Pechstein, N. Tomilin, K. Fredrich, O. Vorontsova, E. Sopova, E. Evergren, V. Haucke, L. Brodin, O. Shupliakov, Vesicle Clustering in a Living Synapse Depends on a Synapsin Region that Mediates Phase Separation, Cell Rep. 30 (2020) 2594–2602.e3. 10.1016/j.celrep.2020.01.092.

[3] C. Hoffmann, J. Rentsch, T.A. Tsunoyama, A. Chhabra, G.A. Perez, R. Chowdhury, F. Trnka, A.A. Korobeinikov, A.H. Shaib, M. Ganzella, G. Giannone, S.O. Rizzoli, A. Kusumi, H. Ewers, D. Milovanovic, Synapsin condensation controls synaptic vesicle sequestering and dynamics, Nat. Commun. 14 (2023) 6730. 10.1038/s41467-023-42372-6.

[4] C. Hoffmann, R. Sansevrino, G. Morabito, C. Logan, R.M. Vabulas, A. Ulusoy, M. Ganzella, D. Milovanovic, Synapsin Condensates Recruit alpha-Synuclein, J. Mol. Biol. (2021) 166961. 10.1016/j.jmb.2021.166961.

[5] R. Sansevrino, C. Hoffmann, D. Milovanovic, Condensate biology of synaptic vesicle clusters, Trends Neurosci. (2023). 10.1016/j.tins.2023.01.001.

[6] D. Park, Y. Wu, S.-E. Lee, G. Kim, S. Jeong, D. Milovanovic, P.D. Camilli, S. Chang, Cooperative function of synaptophysin and synapsin in the generation of synaptic vesicle-like clusters in non-neuronal cells, Nat. Commun. 12 (2021) 263. 10.1038/s41467-020-20462-z.

[7] T.C. Südhof, F. Lottspeich, P. Greengard, E. Mehl, R. Jahn, A synaptic vesicle protein with a novel cytoplasmic domain and four transmembrane regions., Science 238 (1987) 1142 1144.

[8] G. Kim, S.-E. Lee, S. Jeong, J. Lee, D. Park, S. Chang, Multivalent electrostatic pi–cation interaction between synaptophysin and synapsin is responsible for the coacervation, Mol. Brain 14 (2021) 137. 10.1186/s13041-021-00846-y.

[9] F. Gerth, M. Jäpel, A. Pechstein, G. Kochlamazashvili, M. Lehmann, D. Puchkov, F. Onofri, F. Benfenati, A.G. Nikonenko, K. Fredrich, O. Shupliakov, T. Maritzen, C. Freund, V. Haucke, Intersectin associates with synapsin and regulates its nanoscale localization and function., Proc. Natl. Acad. Sci. USA 114 (2017) 12057 12062. 10.1073/pnas.1715341114.

[10] D.S.W. Protter, B.S. Rao, B.V. Treeck, Y. Lin, L. Mizoue, M.K. Rosen, R. Parker, Intrinsically Disordered Regions Can Contribute Promiscuous Interactions to RNP Granule Assembly., Cell Rep 22 (2018) 1401 1412. 10.1016/j.celrep.2018.01.036.

[11] Z. Feng, B. Jia, M. Zhang, Liquid–Liquid Phase Separation in Biology: Specific Stoichiometric Molecular Interactions vs Promiscuous Interactions Mediated by Disordered Sequences, Biochemistry 60 (2021) 2397–2406. 10.1021/acs.biochem.1c00376.

[12] A. Musacchio, On the role of phase separation in the biogenesis of membraneless compartments, EMBO J. 41 (2022) e109952. 10.15252/embj.2021109952.

[13] T.J. Nott, E. Petsalaki, P. Farber, D. Jervis, E. Fussner, A. Plochowietz, T.D. Craggs, D.P. Bazett-Jones, T. Pawson, J.D. Forman-Kay, A.J. Baldwin, Phase transition of a disordered nuage protein generates environmentally responsive membraneless organelles., Mol. Cell 57 (2015) 936 947. 10.1016/j.molcel.2015.01.013.

[14] Y. Lin, S.L. Currie, M.K. Rosen, Intrinsically disordered sequences enable modulation of protein phase separation through distributed tyrosine motifs, J. Biol. Chem. 292 (2017) 19110–19120. 10.1074/jbc.m117.800466.

[15] J. Wang, J.-M. Choi, A.S. Holehouse, H.O. Lee, X. Zhang, M. Jahnel, S. Maharana, R. Lemaitre, A. Pozniakovsky, D. Drechsel, I. Poser, R.V. Pappu, S. Alberti, A.A. Hyman, A Molecular Grammar Governing the Driving Forces for Phase Separation of Prion-like RNA Binding Proteins, Cell 174 (2018) 688–699.e16. 10.1016/j.cell.2018.06.006.

[16] E.W. Martin, A.S. Holehouse, I. Peran, M. Farag, J.J. Incicco, A. Bremer, C.R. Grace, A. Soranno, R.V. Pappu, T. Mittag, Valence and patterning of aromatic residues determine the phase behavior of prion-like domains, Science 367 (2020) 694–699. 10.1126/science.aaw8653.

[17] J.A. Greig, T.A. Nguyen, M. Lee, A.S. Holehouse, A.E. Posey, R.V. Pappu, G. Jedd, Arginine-Enriched Mixed-Charge Domains Provide Cohesion for Nuclear Speckle Condensation, Mol. Cell 77 (2020) 1237–1250.e4. 10.1016/j.molcel.2020.01.025.

[18] K.M. Ruff, Y.H. Choi, D. Cox, A.R. Ormsby, Y. Myung, D.B. Ascher, S.E. Radford, R.V. Pappu, D.M. Hatters, Sequence grammar underlying the unfolding and phase separation of globular proteins, Mol. Cell 82 (2022) 3193–3208.e8. 10.1016/j.molcel.2022.06.024.

[19] A. Patil, A.R. Strom, J.A. Paulo, C.K. Collings, K.M. Ruff, M.K. Shinn, A. Sankar, K.S. Cervantes, T. Wauer, J.D.St. Laurent, G. Xu, L.A. Becker, S.P. Gygi, R.V. Pappu, C.P. Brangwynne, C. Kadoch, A disordered region controls cBAF activity via condensation and partner recruitment, Cell 186 (2023) 4936–4955.e26. 10.1016/j.cell.2023.08.032.

[20] H. Lyons, R.T. Veettil, P. Pradhan, C. Fornero, N.D.L. Cruz, K. Ito, M. Eppert, R.G. Roeder, B.R. Sabari, Functional partitioning of transcriptional regulators by patterned charge blocks, Cell 186 (2023) 327–345.e28. 10.1016/j.cell.2022.12.013.

[21] M.A. Mensah, H. Niskanen, A.P. Magalhaes, S. Basu, M. Kircher, H.L. Sczakiel, A.M.V. Reiter, J. Elsner, P. Meinecke, S. Biskup, B.H.Y. Chung, G. Dombrowsky, C. Eckmann- Scholz, M.P. Hitz, A. Hoischen, P.-M. Holterhus, W. Hülsemann, K. Kahrizi, V.M. Kalscheuer, A. Kan, M. Krumbiegel, I. Kurth, J. Leubner, A.C. Longardt, J.D. Moritz, H. Najmabadi, K. Skipalova, L.S. Blok, A. Tzschach, E. Wiedersberg, M. Zenker, C. Garcia-Cabau, R. Buschow, X. Salvatella, M.L. Kraushar, S. Mundlos, A. Caliebe, M. Spielmann, D. Horn, D. Hnisz, Aberrant phase separation and nucleolar dysfunction in rare genetic diseases, Nature 614 (2023) 564–571. 10.1038/s41586-022-05682-1.

[22] J. Naderi, A.P. Magalhaes, G. Kibar, G. Stik, Y. Zhang, S.D. Mackowiak, H.M. Wieler, F. Rossi, R. Buschow, M. Christou-Kent, M. Alcoverro-Bertran, T. Graf, M. Vingron, D. Hnisz, An activity-specificity trade-off encoded in human transcription factors, Nat. Cell Biol. 26 (2024) 1309–1321. 10.1038/s41556-024-01411-0.

[23] M.R. King, K.M. Ruff, A.Z. Lin, A. Pant, M. Farag, J.M. Lalmansingh, T. Wu, M.J. Fossat, W. Ouyang, M.D. Lew, E. Lundberg, M.D. Vahey, R.V. Pappu, Macromolecular condensation organizes nucleolar sub-phases to set up a pH gradient, Cell 187 (2024) 1889–1906.e24. 10.1016/j.cell.2024.02.029.

[24] J.P. Brady, P.J. Farber, A. Sekhar, Y.-H. Lin, R. Huang, A. Bah, T.J. Nott, H.S. Chan, A.J. Baldwin, J.D. Forman-Kay, L.E. Kay, Structural and hydrodynamic properties of an intrinsically disordered region of a germ cell-specific protein on phase separation, Proc. Natl. Acad. Sci. USA 114 (2017) E8194–E8203. 10.1073/pnas.1706197114.

[25] C.W. Pak, M. Kosno, A.S. Holehouse, S.B. Padrick, A. Mittal, R. Ali, A.A. Yunus, D.R. Liu, R.V. Pappu, M.K. Rosen, Sequence Determinants of Intracellular Phase Separation by Complex Coacervation of a Disordered Protein., Mol. Cell 63 (2016) 72 85. 10.1016/j.molcel.2016.05.042.

[26] I. Alshareedah, W.M. Borcherds, S.R. Cohen, A. Singh, A.E. Posey, M. Farag, A. Bremer, G.W. Strout, D.T. Tomares, R.V. Pappu, T. Mittag, P.R. Banerjee, Sequence-specific interactions determine viscoelasticity and ageing dynamics of protein condensates, Nat. Phys. 20 (2024) 1482–1491. 10.1038/s41567-024-02558-1.

[27] L. Salmon, G. Nodet, V. Ozenne, G. Yin, M.R. Jensen, M. Zweckstetter, M. Blackledge, NMR Characterization of Long-Range Order in Intrinsically Disordered Proteins, J. Am. Chem. Soc. 132 (2010) 8407–8418. 10.1021/ja101645g.

[28] M.R. Jensen, M. Zweckstetter, J. Huang, M. Blackledge, Exploring Free-Energy Landscapes of Intrinsically Disordered Proteins at Atomic Resolution Using NMR Spectroscopy, Chem. Rev. 114 (2014) 6632–6660. 10.1021/cr400688u.

[29] A. Borgia, M.B. Borgia, K. Bugge, V.M. Kissling, P.O. Heidarsson, C.B. Fernandes, A. Sottini, A. Soranno, K.J. Buholzer, D. Nettels, B.B. Kragelund, R.B. Best, B. Schuler, Extreme disorder in an ultrahigh-affinity protein complex, Nature 555 (2018) 61–66. 10.1038/nature25762.

[30] P.O. Heidarsson, D. Mercadante, A. Sottini, D. Nettels, M.B. Borgia, A. Borgia, S. Kilic, B. Fierz, R.B. Best, B. Schuler, Release of linker histone from the nucleosome driven by polyelectrolyte competition with a disordered protein, Nat. Chem. 14 (2022) 224–231. 10.1038/s41557-021-00839-3.

[31] L.G. Randles, S. Batey, A. Steward, J. Clarke, Distinguishing Specific and Nonspecific Interdomain Interactions in Multidomain Proteins, Biophys. J. 94 (2008) 622–628. 10.1529/biophysj.107.119123.

[32] P.E. Wright, H.J. Dyson, Intrinsically disordered proteins in cellular signalling and regulation, Nat. Rev. Mol. Cell Biol. 16 (2015) 18–29. 10.1038/nrm3920.

[33] F. Wiggers, S. Wohl, A. Dubovetskyi, G. Rosenblum, W. Zheng, H. Hofmann, Diffusion of a disordered protein on its folded ligand, Proc. Natl. Acad. Sci. USA 118 (2021) e2106690118. 10.1073/pnas.2106690118.

[34] A.H. Mao, S.L. Crick, A. Vitalis, C.L. Chicoine, R.V. Pappu, Net charge per residue modulates conformational ensembles of intrinsically disordered proteins, Proc. Natl. Acad. Sci. USA 107 (2010) 8183–8188. 10.1073/pnas.0911107107.

[35] R.K. Das, S.L. Crick, R.V. Pappu, N-Terminal Segments Modulate the α-Helical Propensities of the Intrinsically Disordered Basic Regions of bZIP Proteins, J. Mol. Biol. 416 (2012) 287–299. 10.1016/j.jmb.2011.12.043.

[36] R.K. Das, R.V. Pappu, Conformations of intrinsically disordered proteins are influenced by linear sequence distributions of oppositely charged residues, Proc. Natl. Acad. Sci. USA 110 (2013) 13392–13397. 10.1073/pnas.1304749110.

[37] R. van der Lee, M. Buljan, B. Lang, R.J. Weatheritt, G.W. Daughdrill, A.K. Dunker, M. Fuxreiter, J. Gough, J. Gsponer, D.T. Jones, P.M. Kim, R.W. Kriwacki, C.J. Oldfield, R.V. Pappu, P. Tompa, V.N. Uversky, P.E. Wright, M.M. Babu, Classification of intrinsically disordered regions and proteins., Chem. Rev. 114 (2014) 6589 6631. 10.1021/cr400525m.

[38] A.G. Kozlov, E. Weiland, A. Mittal, V. Waldman, E. Antony, N. Fazio, R.V. Pappu, T.M. Lohman, Intrinsically Disordered C-Terminal Tails of E. coli Single-Stranded DNA Binding Protein Regulate Cooperative Binding to Single-Stranded DNA, J. Mol. Biol. 427 (2015) 763– 774. 10.1016/j.jmb.2014.12.020.

[39] R.K. Das, K.M. Ruff, R.V. Pappu, Relating sequence encoded information to form and function of intrinsically disordered proteins, Curr. Opin. Struct. Biol. 32 (2015) 102–112. 10.1016/j.sbi.2015.03.008.

[40] R.K. Das, Y. Huang, A.H. Phillips, R.W. Kriwacki, R.V. Pappu, Cryptic sequence features within the disordered protein p27Kip1 regulate cell cycle signaling, Proc. Natl. Acad. Sci. USA 113 (2016) 5616–5621. 10.1073/pnas.1516277113.

[41] E.W. Martin, A.S. Holehouse, C.R. Grace, A. Hughes, R.V. Pappu, T. Mittag, Sequence Determinants of the Conformational Properties of an Intrinsically Disordered Protein Prior to and upon Multisite Phosphorylation, J. Am. Chem. Soc. 138 (2016) 15323–15335. 10.1021/jacs.6b10272.

[42] T.S. Harmon, M.D. Crabtree, S.L. Shammas, A.E. Posey, J. Clarke, R.V. Pappu, GADIS: Algorithm for designing sequences to achieve target secondary structure profiles of intrinsically disordered proteins, Protein Eng. 29 (2016) 339–346. 10.1093/protein/gzw034.

[43] K.P. Sherry, R.K. Das, R.V. Pappu, D. Barrick, Control of transcriptional activity by design of charge patterning in the intrinsically disordered RAM region of the Notch receptor, Proc. Natl. Acad. Sci. USA 114 (2017) E9243–E9252. 10.1073/pnas.1706083114.

[44] M.K. Shinn, M.C. Cohan, J.L. Bullock, K.M. Ruff, P.A. Levin, R.V. Pappu, Connecting sequence features within the disordered C-terminal linker of Bacillus subtilis FtsZ to functions and bacterial cell division Proc. Natl. Acad. Sci. USA 119 (2022) e2211178119. 10.1073/pnas.2211178119.

[45] M. Borg, T. Mittag, T. Pawson, M. Tyers, J.D. Forman-Kay, H.S. Chan, Polyelectrostatic interactions of disordered ligands suggest a physical basis for ultrasensitivity, Proc. Natl. Acad. Sci. USA 104 (2007) 9650–9655. 10.1073/pnas.0702580104.

[46] J.A. Marsh, J.D. Forman-Kay, Sequence Determinants of Compaction in Intrinsically Disordered Proteins, Biophys. J. 98 (2010) 2383–2390. 10.1016/j.bpj.2010.02.006.

[47] J.D. Forman-Kay, T. Mittag, From Sequence and Forces to Structure, Function, and Evolution of Intrinsically Disordered Proteins, Structure 21 (2013) 1492–1499. 10.1016/j.str.2013.08.001.

[48] T. Zarin, B. Strome, G. Peng, I. Pritišanac, J.D. Forman-Kay, A.M. Moses, Identifying molecular features that are associated with biological function of intrinsically disordered protein regions, Elife 10 (2021) e60220. 10.7554/elife.60220.

[49] S. Müller-Späth, A. Soranno, V. Hirschfeld, H. Hofmann, S. Rüegger, L. Reymond, D. Nettels, B. Schuler, Charge interactions can dominate the dimensions of intrinsically disordered proteins, Proc. Natl. Acad. Sci. USA 107 (2010) 14609–14614. 10.1073/pnas.1001743107.

[50] G. Tesei, A.I. Trolle, N. Jonsson, J. Betz, F.E. Knudsen, F. Pesce, K.E. Johansson, K. Lindorff-Larsen, Conformational ensembles of the human intrinsically disordered proteome, Nature 626 (2024) 897–904. 10.1038/s41586-023-07004-5.

[51] J.M. Lotthammer, G.M. Ginell, D. Griffith, R.J. Emenecker, A.S. Holehouse, Direct prediction of intrinsically disordered protein conformational properties from sequence, Nat. Methods 21 (2024) 465–476. 10.1038/s41592-023-02159-5.

[52] A.S. Holehouse, B.B. Kragelund, The molecular basis for cellular function of intrinsically disordered protein regions, Nat. Rev. Mol. Cell Biol. 25 (2024) 187–211. 10.1038/s41580-023-00673-0.

[53] J.R. Simon, N.J. Carroll, M. Rubinstein, A. Chilkoti, G.P. López, Programming molecular self-assembly of intrinsically disordered proteins containing sequences of low complexity, Nat Chem 9 (2017) 509–515. 10.1038/nchem.2715.

[54] N.S. González-Foutel, J. Glavina, W.M. Borcherds, M. Safranchik, S. Barrera-Vilarmau, A. Sagar, A. Estaña, A. Barozet, N.A. Garrone, G. Fernandez-Ballester, C. Blanes-Mira, I.E. Sánchez, G. de Prat-Gay, J. Cortés, P. Bernadó, R.V. Pappu, A.S. Holehouse, G.W. Daughdrill, L.B. Chemes, Conformational buffering underlies functional selection in intrinsically disordered protein regions, Nat. Struct. Mol. Biol. 29 (2022) 781–790. 10.1038/s41594-022-00811-w.

[55] M.C. Cohan, M.K. Shinn, J.M. Lalmansingh, R.V. Pappu, Uncovering Non-random Binary Patterns Within Sequences of Intrinsically Disordered Proteins, J. Mol. Biol. 434 (2022) 167373. 10.1016/j.jmb.2021.167373.

[56] L. Sawle, K. Ghosh, A theoretical method to compute sequence dependent configurational properties in charged polymers and proteins, J. Chem. Phys. 143 (2015) 085101. 10.1063/1.4929391.

[57] M.V. Staller, E. Ramirez, S.R. Kotha, A.S. Holehouse, R.V. Pappu, B.A. Cohen, Directed mutational scanning reveals a balance between acidic and hydrophobic residues in strong human activation domains, Cell Syst. 13 (2022) 334–345.e5. 10.1016/j.cels.2022.01.002.

[58] I. Langstein-Skora, A. Schmid, R.J. Emenecker, M.O.G. Richardson, M.J. Götz, S.K. Payer, P. Korber, A.S. Holehouse, Sequence- and chemical specificity define the functional landscape of intrinsically disordered regions, BioRxiv (2022) 2022.02.10.480018. 10.1101/2022.02.10.480018.

[59] R.V. Pappu, S.R. Cohen, F. Dar, M. Farag, M. Kar, Phase Transitions of Associative Biomacromolecules, Chem. Rev. 123 (2023) 8945–8987. 10.1021/acs.chemrev.2c00814.

[60] M. Kar, F. Dar, T.J. Welsh, L.T. Vogel, R. Kühnemuth, A. Majumdar, G. Krainer, T.M. Franzmann, S. Alberti, C.A.M. Seidel, T.P.J. Knowles, A.A. Hyman, R.V. Pappu, Phase-separating RNA-binding proteins form heterogeneous distributions of clusters in subsaturated solutions, Proc. Natl. Acad. Sci. USA 119 (2022) e2202222119. 10.1073/pnas.2202222119.

[61] M. Kar, L.T. Vogel, G. Chauhan, S. Felekyan, H. Ausserwöger, T.J. Welsh, F. Dar, A.R. Kamath, T.P.J. Knowles, A.A. Hyman, C.A.M. Seidel, R.V. Pappu, Solutes unmask differences in clustering versus phase separation of FET proteins, Nat. Commun. 15 (2024) 4408. 10.1038/s41467-024-48775-3.

[62] R. Ravindran, I.O.L. Bacellar, X. Castellanos-Girouard, H.M. Wahba, Z. Zhang, J.G. Omichinski, L. Kisley, S.W. Michnick, Peroxisome biogenesis initiated by protein phase separation, Nature 617 (2023) 608–615. 10.1038/s41586-023-06044-1.

[63] M.R. King, K.M. Ruff, R.V. Pappu, Emergent microenvironments of nucleoli, Nucleus 15 (2024) 2319957. 10.1080/19491034.2024.2319957.

[64] Z. Taoufiq, M. Ninov, A. Villar-Briones, H.-Y. Wang, T. Sasaki, M.C. Roy, F. Beauchain, Y. Mori, T. Yoshida, S. Takamori, R. Jahn, T. Takahashi, Hidden proteome of synaptic vesicles in the mammalian brain, Proc. Natl. Acad. Sci. USA 117 (2020) 33586–33596. 10.1073/pnas.2011870117.

[65] A. Denker, K. Kröhnert, J. Bückers, E. Neher, S.O. Rizzoli, The reserve pool of synaptic vesicles acts as a buffer for proteins involved in synaptic vesicle recycling., Proc. Natl. Acad. Sci. USA 108 (2011) 17183 17188. 10.1073/pnas.1112690108.

[66] R.M. Vernon, P.A. Chong, B. Tsang, T.H. Kim, A. Bah, P. Farber, H. Lin, J.D. Forman-Kay, Pi-Pi contacts are an overlooked protein feature relevant to phase separation, Elife 7 (2018) e31486. 10.7554/elife.31486.

[67] R.S. Fisher, S. Elbaum-Garfinkle, Tunable multiphase dynamics of arginine and lysine liquid condensates, Nat. Commun. 11 (2020) 4628. 10.1038/s41467-020-18224-y.

[68] Y. Hong, S. Najafi, T. Casey, J.-E. Shea, S.-I. Han, D.S. Hwang, Hydrophobicity of arginine leads to reentrant liquid-liquid phase separation behaviors of arginine-rich proteins, Nat. Commun. 13 (2022) 7326. 10.1038/s41467-022-35001-1.

[69] M.J. Fossat, X. Zeng, R.V. Pappu, Uncovering Differences in Hydration Free Energies and Structures for Model Compound Mimics of Charged Side Chains of Amino Acids, J. Phys. Chem. B 125 (2021) 4148–4161. 10.1021/acs.jpcb.1c01073.

[70] X. Zeng, K.M. Ruff, R.V. Pappu, Competing interactions give rise to two-state behavior and switch-like transitions in charge-rich intrinsically disordered proteins, Proc. Natl. Acad. Sci. USA 119 (2022) e2200559119. 10.1073/pnas.2200559119.

[71] A. Bremer, M. Farag, W.M. Borcherds, I. Peran, E.W. Martin, R.V. Pappu, T. Mittag, Deciphering how naturally occurring sequence features impact the phase behaviours of disordered prion-like domains, Nat. Chem. 14 (2022) 196–207. 10.1038/s41557-021-00840-w.

[72] S.-H. Oh, J. Lee, M. Lee, S. Kim, W.B. Lee, D.W. Lee, S.-H. Choi, Simple Coacervation of Guanidinium-Containing Polymers Induced by Monovalent Salt, Macromolecules 56 (2023) 3989–3999. 10.1021/acs.macromol.2c02346.

[73] P.L. Cameron, T.C. Südhof, R. Jahn, P.D. Camilli, Colocalization of synaptophysin with transferrin receptors: implications for synaptic vesicle biogenesis., J. Cell Biol. 115 (1991) 151–164. 10.1083/jcb.115.1.151.

[74] W.E. Arter, R. Qi, N.A. Erkamp, G. Krainer, K. Didi, T.J. Welsh, J. Acker, J. Nixon-Abell, S. Qamar, J. Guillén-Boixet, T.M. Franzmann, D. Kuster, A.A. Hyman, A. Borodavka, P.S. George-Hyslop, S. Alberti, T.P.J. Knowles, Biomolecular condensate phase diagrams with a combinatorial microdroplet platform, Nat. Commun. 13 (2022) 7845. 10.1038/s41467-022-35265-7.

[75] A. Chattaraj, M.L. Blinov, L.M. Loew, The solubility product extends the buffering concept to heterotypic biomolecular condensates, Elife 10 (2021) e67176. 10.7554/elife.67176.

[76] M. Farag, W.M. Borcherds, A. Bremer, T. Mittag, R.V. Pappu, Phase separation of protein mixtures is driven by the interplay of homotypic and heterotypic interactions, Nat. Commun. 14 (2023) 5527. 10.1038/s41467-023-41274-x.

[77] D. Qian, H. Ausserwoger, T. Sneideris, M. Farag, R.V. Pappu, T.P.J. Knowles, Dominance analysis to assess solute contributions to multicomponent phase equilibria, Proc. Natl. Acad. Sci. USA 121 (2024) e2407453121. 10.1073/pnas.2407453121.

[78] A.N. Semenov, M. Rubinstein, Thermoreversible Gelation in Solutions of Associative Polymers. 1. Statics, Macromolecules 31 (1998) 1373–1385. 10.1021/ma970616h.

[79] J.-M. Choi, A.A. Hyman, R.V. Pappu, Generalized models for bond percolation transitions of associative polymers, Phys. Rev. E 102 (2020) 042403. 10.1103/physreve.102.042403.

[80] F. Tanaka, Molecular Gels, (2006) 17–77. 10.1007/1-4020-3689-2_2.

[81] M. Rubinstein, A.N. Semenov, Thermoreversible Gelation in Solutions of Associating Polymers. 2. Linear Dynamics, Macromolecules 31 (1998) 1386–1397. 10.1021/ma970617+.

[82] T.S. Harmon, A.S. Holehouse, M.K. Rosen, R.V. Pappu, Intrinsically disordered linkers determine the interplay between phase separation and gelation in multivalent proteins, Elife 6 (2017) e30294. 10.7554/elife.30294.

[83] P.J. Flory, Thermodynamics of High Polymer Solutions, J. Chem. Phys. 10 (1942) 51–61. 10.1063/1.1723621.

[84] M.L. Huggins, Solutions of Long Chain Compounds, J. Chem. Phys. 9 (1941) 440–440. 10.1063/1.1750930.

[85] Y. Song, J. Zhang, D. Li, Microfluidic and Nanofluidic Resistive Pulse Sensing: A Review, Micromachines 8 (2017) 204. 10.3390/mi8070204.

[86] R.W. DeBlois, C.P. Bean, Counting and Sizing of Submicron Particles by the Resistive Pulse Technique, Rev. Sci. Instrum. 41 (1970) 909–916. 10.1063/1.1684724.

[87] S.L. Crick, K.M. Ruff, K. Garai, C. Frieden, R.V. Pappu, Unmasking the roles of N- and C-terminal flanking sequences from exon 1 of huntingtin as modulators of polyglutamine aggregation, Proc. Natl. Acad. Sci. USA 110 (2013) 20075–20080. 10.1073/pnas.1320626110.

[88] M. Kar, A.E. Posey, F. Dar, A.A. Hyman, R.V. Pappu, Glycine-Rich Peptides from FUS Have an Intrinsic Ability to Self-Assemble into Fibers and Networked Fibrils, Biochemistry 60 (2021) 3213–3222. 10.1021/acs.biochem.1c00501.

[89] C. Hoffmann, G. Murastov, J.V. Tromm, J.-B. Moog, M.A. Aslam, A. Matkovic, D. Milovanovic, Electric Potential at the Interface of Membraneless Organelles Gauged by Graphene, Nano Lett. (2023). 10.1021/acs.nanolett.3c02915.

[90] Z. Farsi, J. Preobraschenski, G. van den Bogaart, D. Riedel, R. Jahn, A. Woehler, Single-vesicle imaging reveals different transport mechanisms between glutamatergic and GABAergic vesicles, Science 351 (2016) 981–984. 10.1126/science.aad8142.

[91] E. Kosmidis, C.G. Shuttle, J. Preobraschenski, M. Ganzella, P.J. Johnson, S. Veshaguri, J. Holmkvist, M.P. Møller, O. Marantos, F. Marcoline, M. Grabe, J.L. Pedersen, R. Jahn, D. Stamou, Regulation of the mammalian-brain V-ATPase through ultraslow mode-switching, Nature 611 (2022) 827–834. 10.1038/s41586-022-05472-9.

[92] P. Li, S. Banjade, H.-C. Cheng, S. Kim, B. Chen, L. Guo, M. Llaguno, J.V. Hollingsworth, D.S. King, S.F. Banani, P.S. Russo, Q.-X. Jiang, B.T. Nixon, M.K. Rosen, Phase transitions in the assembly of multivalent signalling proteins., Nature 483 (2012) 336 340. 10.1038/nature10879.

[93] A.C. Kokotos, C.B. Harper, J.R.K. Marland, K.J. Smillie, M.A. Cousin, S.L. Gordon, Synaptophysin sustains presynaptic performance by preserving vesicular synaptobrevin-II levels, J. Neurochem. 151 (2019) 28–37. 10.1111/jnc.14797.

[94] S.L. Gordon, R.E. Leube, M.A. Cousin, Synaptophysin Is Required for Synaptobrevin Retrieval during Synaptic Vesicle Endocytosis, J. Neurosci. 31 (2011) 14032–14036. 10.1523/jneurosci.3162-11.2011.

[95] M. Atias, Y. Tevet, J. Sun, A. Stavsky, S. Tal, J. Kahn, S. Roy, D. Gitler, Synapsins regulate α-synuclein functions., Proc. Natl. Acad. Sci. USA 116 (2019) 11116 11118. 10.1073/pnas.1903054116.

[96] J. Sun, L. Wang, H. Bao, S. Premi, U. Das, E.R. Chapman, S. Roy, Functional cooperation of α-synuclein and VAMP2 in synaptic vesicle recycling., Proc. Natl. Acad. Sci. USA 116 (2019) 11113–11115. 10.1073/pnas.1903049116.

[97] X.M. Zhang, U. François, K. Silm, M.F. Angelo, M.V. Fernandez-Busch, M. Maged, C. Martin, V. Bernard, F.P. Cordelières, M. Deshors, S. Pons, U. Maskos, A.P. Bemelmans, S.M. Wojcik, S.E. Mestikawy, Y. Humeau, E. Herzog, A proline-rich motif on VGLUT1 reduces synaptic vesicle super-pool and spontaneous release frequency, Elife 8 (2019) e50401. 10.7554/elife.50401.

[98] J. Jackson, C. Hoffmann, E. Scifo, H. Wang, L. Wischhof, A. Piazzesi, M. Mondal, H. Shields, X. Zhou, M. Mondin, E.B. Ryan, H. Döring, J.H.M. Prehn, K. Rottner, G. Giannone, P. Nicotera, D. Ehninger, D. Milovanovic, D. Bano, Actin-nucleation promoting factor N-WASP influences alpha-synuclein condensates and pathology, Cell Death Dis. 15 (2024) 304. 10.1038/s41419-024-06686-7.

[99] D. Park, K. Fujise, Y. Wu, R. Luján, S.D. Olmo-Cabrera, J.F. Wesseling, P.D. Camilli, Overlapping role of synaptophysin and synaptogyrin family proteins in determining the small size of synaptic vesicles, Proc. Natl. Acad. Sci. USA 121 (2024) e2409605121. 10.1073/pnas.2409605121.

[100] T.U. Consortium, UniProt: the Universal Protein Knowledgebase in 2023, Nucleic Acids Res. 51 (2022) D523–D531. 10.1093/nar/gkac1052.

[101] D. Piovesan, A.D. Conte, D. Clementel, A.M. Monzon, M. Bevilacqua, M.C. Aspromonte, J.A. Iserte, F.E. Orti, C. Marino-Buslje, S.C.E. Tosatto, MobiDB: 10 years of intrinsically disordered proteins, Nucleic Acids Res. 51 (2022) D438–D444. 10.1093/nar/gkac1065.

[102] A. Hubstenberger, M. Courel, M. Bénard, S. Souquere, M. Ernoult-Lange, R. Chouaib, Z. Yi, J.-B. Morlot, A. Munier, M. Fradet, M. Daunesse, E. Bertrand, G. Pierron, J. Mozziconacci, M. Kress, D. Weil, P-Body Purification Reveals the Condensation of Repressed mRNA Regulons, Mol. Cell 68 (2017) 144–157.e5. 10.1016/j.molcel.2017.09.003.

[103] S. Jain, J.R. Wheeler, R.W. Walters, A. Agrawal, A. Barsic, R. Parker, ATPase-Modulated Stress Granules Contain a Diverse Proteome and Substructure, Cell 164 (2016) 487–498. 10.1016/j.cell.2015.12.038.

[104] J.H. Ward, Hierarchical Grouping to Optimize an Objective Function, J. Am. Stat. Assoc. 58 (1963) 236. 10.2307/2282967.

[105] J. Huerta-Cepas, D. Szklarczyk, D. Heller, A. Hernández-Plaza, S.K. Forslund, H. Cook, D.R. Mende, I. Letunic, T. Rattei, L.J. Jensen, C. von Mering, P. Bork, eggNOG 5.0: a hierarchical, functionally and phylogenetically annotated orthology resource based on 5090 organisms and 2502 viruses, Nucleic Acids Res. 47 (2019) D309–D314. 10.1093/nar/gky1085.

[106] S.F. Altschul, W. Gish, W. Miller, E.W. Myers, D.J. Lipman, Basic local alignment search tool, J. Mol. Biol. 215 (1990) 403–410. 10.1016/s0022-2836(05)80360

[107] R.C. Edgar, MUSCLE: multiple sequence alignment with high accuracy and high throughput, Nucleic Acids Res. 32 (2004) 1792–1797. 10.1093/nar/gkh340.

[108] A.M. Waterhouse, J.B. Procter, D.M.A. Martin, M. Clamp, G.J. Barton, Jalview Version 2—a multiple sequence alignment editor and analysis workbench, Bioinformatics 25 (2009) 1189–1191. 10.1093/bioinformatics/btp033.

[109] C. Akshita, H. Christian, K.A. Aleksandr, R. Jakob, K. Linda, G. Luka, R.-V. Cristina, J.C. Emma, W.N. Jaqulin, R. Branislava, P. Eleonora, K. Sarah, R.O. Silvio, E. Helge, M.R. Jennifer, M. Dragomir, Condensates of synaptic vesicles and synapsin are molecular beacons for actin sequestering and polymerization, BioRxiv (2024) 2024.07.19.604346. 10.1101/2024.07.19.604346.

[110] D. Qin, Y. Xia, G.M. Whitesides, Soft lithography for micro- and nanoscale patterning., Nat. Protoc. 5 (2010) 491–502. 10.1038/nprot.2009.234.

[111] N.A. Erkamp, T. Sneideris, H. Ausserwöger, D. Qian, S. Qamar, J. Nixon-Abell, P.S. George-Hyslop, J.D. Schmit, D.A. Weitz, T.P.J. Knowles, Spatially non-uniform condensates emerge from dynamically arrested phase separation, Nat. Commun. 14 (2023) 684. 10.1038/s41467-023-36059-1.

[112] H. Ausserwöger, R. Scrutton, T. Sneideris, C.M. Fischer, D. Qian, E. de Csilléry, K.L. Saar, A.Z. Białek, M. Oeller, G. Krainer, T.M. Franzmann, S. Wittmann, J.M. Iglesias-Artola, G. Invernizzi, A.A. Hyman, S. Alberti, N. Lorenzen, T.P.J. Knowles, Biomolecular condensates sustain pH gradients at equilibrium driven by charge neutralisation, BioRxiv (2024) 2024.05.23.595321. 10.1101/2024.05.23.595321.

